# High-Speed Mass Spectrometers diminish the difference between Data-Dependent and Data-Independent Acquisition Proteomics

**DOI:** 10.64898/2026.05.26.727836

**Authors:** Naomi O’Sullivan, Florian Bayer, Carolin Mogler, Bernhard Kuster

## Abstract

Data-dependent acquisition mass spectrometry (DDA-MS) and data-independent acquisition mass spectrometry (DIA-MS) have historically offered complementary strengths in bottom-up proteomics, with DDA providing high-selectivity spectra for post-translational modification (PTM) analysis and DIA enabling more systematic peptide sampling. Here, we asked if this is still the case for the Orbitrap Astral platform that offers high-speed DDA and (ultra-) narrow-window DIA (nDIA) capabilities across proteome and phosphoproteome applications. When DDA and DIA measurements were parameter-matched (to the extent possible), the differences in analytical performance diminished markedly. Across extensive replicate analyses, both methods continued to identify new peptides and proteins without reaching saturation, indicating that the molecular complexity of biological samples still overwhelms even the fastest liquid chromatography-MS (LC-MS) methods. Incomplete sampling also contributed to substantial peptide-level non-overlap between DDA and nDIA and data completeness was only modestly better for nDIA than DDA across many replicates. Quantitatively, DDA and nDIA showed broadly similar precision and accuracy, with nDIA offering slightly higher precision and DDA slightly better accuracy in controlled mixture experiments. MS1-based quantification outperformed MS2-based quantification, particularly for short gradients, supporting MS1 quantification as a robust and general strategy for high-throughput proteomics. In phosphoproteomic samples, DDA and nDIA identified similar numbers of phosphopeptides, but DDA retained a small edge for phosphorylation site localisation. Together, the results show that advances in acquisition speed and sensitivity are narrowing the historical gap between DDA and DIA, while also revealing that current LC-MS workflows remain far from providing comprehensive proteome coverage. Going forward, further gains in dynamic range, scan speed, sensitivity, and transparent software tools will be required to reach systematic, comprehensive and reliable measurements of complex proteomes in a single shot.

## Introduction

Liquid chromatography (LC) coupled to data-dependent acquisition mass spectrometry (DDA-MS) has paved the way for the remarkable development of the field of proteomics. It followed the simple logic that the vast molecular complexity of trypsin-digested proteomes demands very high separation power and sensitivity of the analytical system to identify unambiguously and quantify robustly individual constituent components. Beyond important contributions of the LC part of an LC-MS system, DDA achieved both by i) acquiring mass spectra of peptides (MS1) at high resolution and mass accuracy; ii) selecting a very narrow window (e. g. 1.2 Th) of the mass to charge ratio (m/z) range of tryptic peptides (300-2,000) for fragmentation; iii) controlling the collision energy for individual peptides and iv) acquiring low-complexity fragment ion spectra of fragment ions (MS2). Together with sophisticated software for database searching and artificial intelligence-based rescoring at low false discovery rates (FDR)^1^, these analytical characteristics have supported countless small and large-scale proteome projects. In DDA, peptide quantification uses the MS1 signal of intact peptides. To work robustly, this requires adapting the cycle time of DDA-MS measurements (i. e. the time between acquiring subsequent MS1 spectra) to the peak widths provided by the LC separation. Ideally, 8-10 MS1 spectra are collected for any peptide across its LC peak. DDA parameters also allow control over collecting peptide ions for controllable lengths of time to tune sensitivity. The very high selectivity of DDA-MS has been instrumental for the confident analysis of post-translational modifications (PTMs), open modification searches for discovering new PTMs^2^ as well as de-novo peptide sequencing^3^. However, the semi-stochastic nature of selecting peptides for fragmentation, typically based on intensity and charge, and the finite signal to noise (SN) ratio with which (low abundance) peptides can be detected in MS1, results in only partial coverage of peptides eluting from the LC at any one time and in any one analysis. Therefore, the overlap (or data completeness) between analyses is often modest. While both improve with increasing LC separation time as well as the speed and sensitivity of the mass spectrometer DDA cannot achieve systematic and complete coverage of complex proteomes.

Data-independent acquisition mass spectrometry (DIA-MS) offers a conceptually powerful alternative. While it shares many of the features of DDA mentioned above, DIA systematically measures (ideally) all peptides eluting from the LC at any one time across the chosen MS1 m/z range. This increases peptide coverage as well as data completeness between analyses of many samples. One limitation of DIA has always been the need to balance the resolution of the LC separation with the speed at which the mass spectrometer can collect data to fulfil the requirement of collecting 8-10 MS1 or MS2 spectra for robust quantification. In other words, the more performant the LC system or the shorter the LC gradients, the higher the duty cycle of the MS must be. In fact, the earliest implementation of DIA-MS did not mass-select peptide ions for fragmentation, but instead operated the MS instrument in alternating cycles of low-energy (to collect peptide precursor data) and high-energy (to collect fragment ion data) collision-induced dissociation (CID) mode^4^. This very high duty cycle came at the expense of collecting extremely complex fragment ion mass spectra which were difficult, to impossible, to deconvolute into individual peptides.

Many DIA scan modes emerged that sought to find good compromises between the somewhat incompatible goals of collecting data systematically and consistently for all peptides in all samples and collecting this data quickly to enable large-scale applications. Important parameters to consider (many but not all shared with DDA) in this context are: i) the (narrow) peak widths of peptides separated by high-performance LC systems, ii) the (high) dynamic abundance range of peptide eluting at the same time, iii) the (broad) range of mass-to-charge ratios (m/z) that the mass spectrometer has to cover, iv) the (different) optimal collision energies of peptides for fragmentation, vi) the (slow) duty cycle of the mass spectrometer and v) the ability of software to deconvolute (complex) peptide fragment ion spectra. A common feature of all current DIA-MS modes is that they use some form of peptide ion selection prior to fragmentation to reduce MS2 spectral complexity. For instance, the SWATH-MS^5,6^ approach employed quadrupole selection of peptides in fixed or variable “windows” of typically 10-100 Da to cover the desired m/z range^7^. A variation on this theme is scanning DIA^8,9^ which achieves higher ion selectivity by continuously sweeping the isolation window across the chosen m/z range. In addition, ion mobility spectrometry permits on-the-fly ion fractionation prior to peptide fragmentation^10–12^. As for DDA-MS, many of the compromises mentioned above become lesser issues with increasing the speed and sensitivity of mass spectrometers but it is currently unclear e. g. “how fast is fast enough” for DIA-MS.

The recent introduction of very fast (100-300 Hz) and sensitive (single ion detection) hybrid mass spectrometers such as the Orbitrap^13,14^, timsTOF^10^, ZenoTOF^15^ and Xevo MRT platforms marked a step change. They feature elements of ion accumulation, ion selection, time-of-flight ion separation, and single-ion detection, which collectively enhance sensitivity, reduce spectral complexity, and increase speed. These hardware improvements have enabled the implementation of narrow-window DIA^16^ (nDIA) that isolates peptides with a 2-4 Th window, thus approaching the selection window of typical DDA measurements. While DIA and nDIA have recently been benchmarked for different applications^16–21^, the potential of DDA on this platform and to what extent nDIA and DDA measurements become the same has not been systematically explored^16,22^. The results presented in this study indicate that the differences between DDA and DIA are indeed diminishing. Briefly, DIA consistently led to higher peptide and protein identifications in single analysis, but peptidome and proteome saturation could not be achieved by both methods even when analysing many replicates. Controlled two species mixture analysis showed that quantitative precision was slightly higher for DIA, but DDA was slightly more accurate. MS1 quantification for DIA turned out to be superior to MS2 quantification particularly when running short LC gradients and using ultra narrow isolation windows. For phosphopeptides, DDA and nDIA identified very similar numbers but DDA outperformed DIA for site localisation.

Discrepancies that could not be fully explained included: i) the fact that a similar number and substantial fraction of peptides (28%) could only be found by either DDA or DIA and ii) that data completeness was only marginally higher for DIA when analysing many samples. Both suggest that: i) the molecular complexity of proteome-scale samples still overwhelms the LC-MS system so that there are more stochastic elements at work in the LC-MS/MS workflow than generally appreciated and ii) the assumptions made by software algorithms used for the combined analysis of larger sets of DDA and DIA data have a larger influence on results than they should, an issue that the proteomics community needs to address^23,24^.

## Experimental Procedures

### Experimental Design and Statistical Rationale

To provide a robust technical evaluation, sample types were limited to HeLa (all figures) and *Escherichia coli* (Fig. 3) peptides. Measurements across different methods were taken using the same peptide pool to ensure sample equivalency, often drawing directly from the same vial. To further minimise potential bias, notably from regards to mass or retention time (RT) drift, Data-Dependant Acquisition (DDA) and Data-Independent Acquisition (DIA) measurements were acquired in an interleaved manner across methods. This strategy ensured both methods are equally affected by technical variability. The majority of experiments were acquired in technical triplicates to perform standard deviation and coefficients of variation calculations, to explore identification and quantification repeatability. No biological replicates were acquired as it was not relevant to this study. DDA parameter exploration (Supplemental Fig. 1) was performed without replicates (n=1) as these were exploratory measurements intended to identify general trends rather than make statistical inferences. For Supplemental Figure 2, thirty replicates (n=30) were acquired to reduce stochastic sampling variability and achieve robust characterization of peptide properties. Data analysis was primarily descriptive, focusing on consistent trends and large effect sizes observed between acquisition methods rather than formal statistical hypothesis testing. For Fig. 1D, a paired student’s t-test was performed.

**Figure 1.**
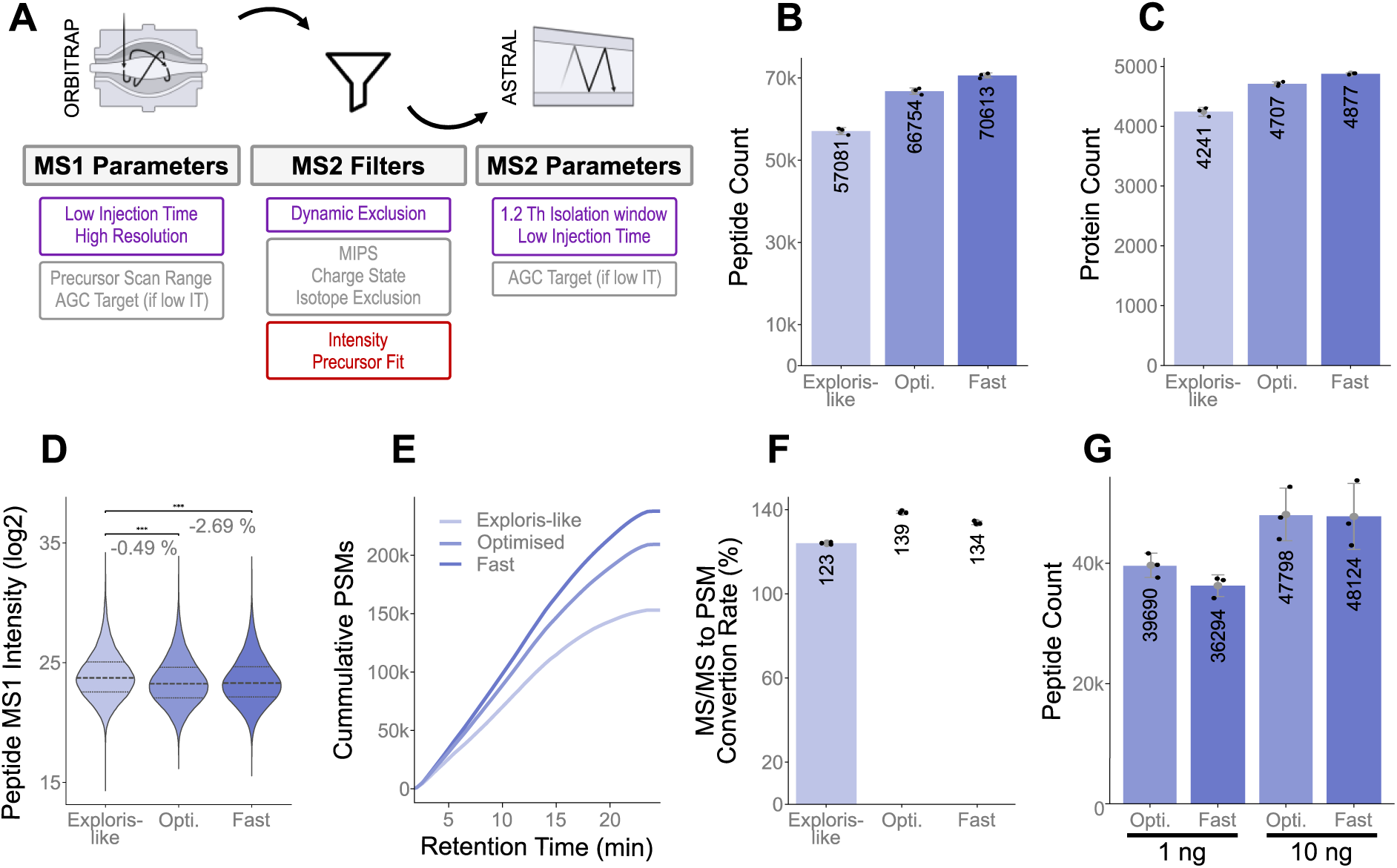
Comparison of Orbitrap DDA acquisition methods combining optimised parameters and scan speed. (A) Schematic overview of the MS1 and MS2 parameters affecting Orbitrap Astral performance, summarising positive (purple), neutral (grey) and negative (red) effects on peptide identifications. (B) Bar plot showing the number of peptide identifications (mean +/- standard deviation, n=3) for three methods: an “Exploris-like” reference method, an “Optimised” method implementing best parameter combinations, and a “Fast” method designed to maximize acquisition speed. (C) Same as panel (B) but for protein identifications. Protein identifications required two or more peptides and a minimum of one unique peptide. (D) Violin plots illustrating the distribution of (log2) peptide intensities for each method. Medians, upper and lower quartiles are marked with dashed lines. Statistical comparisons between paired conditions were performed using a two-tailed paired Student’s t-test. *** indicates p<0.001. (E) Line plot showing the cumulative number of identified peptide-spectrum matches (PSM) over the chromatographic gradient for each of the three methods. (F) Bar plot showing the ratio (in %) of PSMs over the number of acquired MS2 spectra for the three methods (conversion rate, mean +/- standard deviation, n=3). (G) Bar plot showing the number of identified peptides (peptide count) for two methods and two sample loadings (mean +/- standard deviation, n=3).

### Full Proteome Peptide Preparation

HeLa cells were cultured in Dulbecco’s Modified Eagle Medium (DMEM; Gibco, Invitrogen), supplemented with 10% fetal bovine serum, at 37 °C, in a humidified incubator (5% CO_2_). Cells were harvested at ∼80% confluence by washing twice with Phosphate-buffered saline (PBS) buffer and lysed in 8 M urea, 40 mM tris(hydroxymethyl)aminomethane-hydrochloride (Tris/HCl, pH 7.6), 1 × ethylenediaminetetraacetic acid (EDTA)-free protease inhibitor mixture (Complete Mini, Roche). The cell lysate was clarified by centrifuged at 20,000 × g for 20 min at 4°C. Protein concentration was determined through Bicinchoninic acid (BCA) protein assay (Pierce). Proteins were reduced using 10mM dithiothreitol (DTT) and incubated 30 min at 30°C and 700rpm on a thermoshaker. Cysteines were alkylated using 50mM chloroacetamide (CAA) and incubated at room temperature for 30 minutes in the dark. Samples were diluted 1:5 using 40mM Tris/HCl to reach a urea concentration < 1.6M. Proteins were pre-digested with Lys-C in a protease-to-protein ratio of 1:100 (w/w) for 4 hours, at 37°C and 600 rpm. Trypsin (Roche) was added in a protease-to-protein ratio of 1:100 (w/w) and incubated overnight in the same conditions. Protease activity was halted by the addition of formic acid (FA) (1% v/v). The tryptic peptides were desalted using SepPak 200mg cartridges according to the user manual. The cartridges are washed once with 1 mL acetonitrile (ACN), once with 1 mL 0.1% FA in 50% ACN, then twice with 1 mL 0.1% FA in water. The acidified samples are loaded onto the column and the flowthrough collected. The flowthrough was reloaded and the column washed twice with 1 mL 0.1% FA. The peptides were eluted using 2 x 200 uL 0.1% FA in 50% ACN. Eluted peptides were either stored at - 80°C or vacuum dried before mass spectrometry analysis.

### Phospho-Proteome Peptide Preparation

Hela cells were cultured as described above and lysed in 2% sodium dodecyl sulphate (SDS), 10 mM Tris/HCl (pH 7.6). The cell lysate was heated 10 min at 95°C on a thermoshaker at 400 rpm. Trifluoroacetic acid (TFA) was added for a final concentration of 1% (v/v), briefly mixed then N-Methylmorpholine (NMM) was added to reach a final concentration of 2% (∼pH 7-8). Protein concentration was determined through BCA protein assay (Pierce). Proteins were digested according to the Single-Pot, Solid-Phase, Sample-Preparation (SP3) protocol on an Agilent Bravo handling platform^25^. In brief, 20 μl Sera Mag A–B bead mix (1:1) was used per 200 μg total protein lysate sample and proteins were bound at a final concentration of 70% ethanol. Beads were washed with 80% ethanol and acetonitrile. Proteins were reduced using 10mM tris(2-carboxyethyl)phosphine (TCEP) and cysteines alkylated using 50 mM CAA for 30 minutes at 37°C in the dark at 800 rpm. Bead-bound proteins were digested with trypsin (Roche) in a protease-to-protein ratio of 1:100 (w/w) and incubated overnight at 37°C on a thermoshaker at 800 rpm. The next day, peptides were collected, the beads washed in 0.1% FA, then acidified sufficiently to reach 1% FA (v/v). The peptides were desalted using SepPak 200mg cartridges as described above and vacuum dried. Phospho-peptide enrichment was performed by immobilized metal ion affinity chromatography (IMAC) enrichment on a ProPac 0.5 ml Fe(III)-loaded NTA column (Thermo Scientific), according to Ruprecht et.al^26^. Using an Aekta Pure Liquid-Chromatography (LC) system (GE Healthcare Life Sciences), samples were manually injected into the loop, then loaded on the column with a flow rate of 0.2 ml/min. Bound phosphopeptides were eluted and collected applying a two-step gradient from 0 % to 12 % IMAC elution solvent (0.315 % NH4OH) in 1.5 min at 0.55 ml/min and to 26 % elution solvent in 5 min at 3 ml/min. The phospho-peptide peak was manually collected and acidified with TFA to a final concentration of 1 % TFA. Eluted peptides were dried by vacuum centrifugation and cleaned using SepPak cartridges.

### *Escherichia coli* Peptide and Species Mix Preparation

*Escherichia coli* samples were prepared as in Abele et al.^27,28^. Briefly, *E. Coli* cell pellets were lysed with 100% TFA, incubated at room temperature for 5 minutes before being neutralised in 10x 2M Tris volume. Proteins were reduced and alkylated using 9 mM TCEP and 40mM CAA before sample dilution (<5% TFA and <1M Tris) and overnight digestion using trypsin-to-protein ratio of 1:50 (w/w). Tryptic peptides were desalted using self-packed tips with Empore C18 disks. Tips were primed with 100% ACN, then 40% ACN/0.1% FA, and finally equilibrated with 0.1% FA. Peptides were loaded, washed with 0.1% FA, and eluted twice with 0.1% FA in 40% ACN. The *E. Coli* peptides were mixed into HeLa peptides at 1:2 or 2:1 ratio.

### Liquid Chromatography Separation

Samples were analysed using the Vanquish Neo UHPLC system (Thermo Fisher Scientific) coupled to an Orbitrap Astral mass spectrometer (Thermo Fisher Scientific) with a Nanospray Flex source.

The LC system was operated in trap and elute mode using trap backwards flush. Peptides were loaded onto a PepMap Neo Trap Cartridge (300 µm x 5 mm) and then separated using an Ionopticks Aurora Rapid (75 µm x 8 cm, Ionopticks) analytical column at 50°C. Mobile phase A was 0.1% FA in water and mobile phase B, 0.1% FA in ACN. The majority acquisitions were made using the following gradient: 2% to 6% B in 0.5 minutes with a flow rate of 750nl/min, 6% to 8% B in 2 minutes while decreasing the flow rate to 300nl/min, 8% to 32% for 19 minutes, column washing at 80% B at 750nl/min for 2 minutes and column equilibration at 2% B using flow rate and pressure combined control (column equilibration factor 3). The trap cartridge is washed in parallel using 4 zebra wash cycles.

For quantitatively matched methods (Fig. 4), different active gradient lengths were used: 5-, 15-, 30- and 60-minutes. The flow rate was set to 300nl/min throughout the active gradient and increased to 750nl/min for column washing and equilibration. Washing and equilibration was carried out as described above. The active gradients were as follows: mobile phase B was increased from 2% to 6% in 0.2, 0.5 and 0.5 minutes, then increased to 32% B for a duration of 4.8, 14.5 or 29.5 minutes to reach a total of 5-, 15- or 30-minutes respectively. For the 60-minute gradient, solvent B was increased from 1% to 28% linearly.

Finally, a limited number of acquisitions used to illustrate some DDA parameter optimisations (Supplemental Fig. 1) used an in-house packed column (75 um x 48cm; 3 um C18 resin, Reprosil PUR AQ - Dr. Maisch). Separation was achieved using 2%-4% B in 1 min and 4%-32% over 27 min followed by the column wash at 80% B for 3 min.

### Mass Spectrometry Analysis

Given the extensive parameter testing performed for initial DDA method development then multiple method comparisons, a comprehensive summary of parameters used is provided in Supplemental Table 1. For the initial DDA experiments, three acquisition methods were established: ‘Exploris-like’, ‘Optimised’ and ‘Fast’. For the ‘Exploris-like’ method, MS1 full scans were recorded using the Orbitrap mass analyser at a resolution of 120000 every 1 second. The Automatic Gain Control (AGC) target was set to 300% (3e6 charges), the maximal injection time (maxIT) to 50 ms and the precursor scan range as 360-1300 m/z. In parallel, MS/MS spectra were acquired in the Astral mass analyser. MS/MS spectra were acquired using a normalised collision energy (NCE) of 26%, a 1.2 Th isolation window, an AGC target of 300% (3e4 charges) and a maxIT of 10 ms across a 100-2000 m/z scan range. The Monoisotopic Precursor Selection (MIPS) filter mode was set to peptide, with restriction relaxed if too few precursors were available. Only precursors of charge states 2-6 were selected, as well as undetermined charge states. The dynamic exclusion filter was used to exclude precursors for 30 seconds after picking within a 10 ppm tolerance, while also excluding isotopes. A minimal intensity filter was also applied for a minimum of 1e4.

For the Optimised and Fast methods, MS1 full scans were acquired using a resolution of 240000 every 0.6 seconds. MS1 maxIT was set to either 50 ms or 3 ms and MS2 maxIT to 3.5ms or 5ms for the optimised and fast methods respectively. Precursor scan range was set to 360-1300 m/z for the optimisation method but was slightly reduced to 360-1100 m/z for the fast method. Dynamic exclusion only excluded precursors for 5 seconds after picking within a 5 ppm tolerance and no intensity filter was applied. For subsequent experiments, the ‘fast’ method was utilised with minor changes: all quantitatively matched methods had a shorter scan range (360-1000 m/z) and for the 5 minute measurement, a lower resolution and faster cycle time was used (180000, 0.5sec); phosphopeptides were measured with a wider scan range (360-1300 m/z). All other parameters were the same.

For DIA methods, MS1 full scans were recorded with a resolution of 240000 every 0.6 seconds. The AGC target was set to 300%, the maxIT to 3 ms and the precursor scan range as 360-1000 m/z. Ions were fragmented using an NCE of 28%. nDIA MS/MS spectra were then acquired using an AGC target of 300% (3e4 charges) and a maximal IT of 3 ms across a 150-2000 m/z scan range. Unless specified, MS/MS acquisition used a 2 Th isolation window; 1.2 Th and 4 Th windows were applied where indicated. To achieve the quantitatively matched methods, scan ranges and isolation windows were adjusted to the gradient length for sufficient peak sampling. For a fixed scan range (360-1000 m/z), window sizes were set to 6.5 Th, 3.3 Th, 2.2 Th and 1.2 Th for 5-, 15-, 30- and 60-minute gradients respectively.

Alternatively, for methods employing a fixed window size (2 Th), the scan ranges were adjusted to 580-750 m/z, 490-870 m/z, 400-960 m/z and 350-1350 m/z to match gradient lengths. Additional details are provided in the main text and Supplemental Table 1.

### Search rationale

FragPipe was chosen for data processing due to its capacity to handle both DDA and DIA data and the similarities and flexibility in the processing frameworks. The decreased spectral complexity afforded by narrow window DIA limits the spectral deconvolution required by DIA-processing algorithms, resulting in more DDA-like spectra. In opposition, the development of DDA algorithms accepting chimeric spectra^29,30^ applies DIA-like spectral deconvolution to DDA spectra. This suggests that many aspects of data processing could be harmonized across acquisition methods. FragPipe utilizes many common modules for proteomic searches and has described using similar strategies for both DDA+ and DIA analysis through the different MSFragger mode^29,31^. As described in Yu et al., DIA spectra are searched against all database peptides within the possible precursor mass range, providing a list of candidate peptides for which peaks and features are subsequently determined. The peptide candidate list can be filtered and rescored before selecting the top peptide spectrum match (PSM) (rank1) for the associated spectrum. Due to the chimeric nature of DIA data, the matched fragments are subtracted from the spectrum and the members of the candidate list rescored to obtain the second ranked PSM (rank 2).

This is repeated until too few peaks or candidates remain. The DDA+ strategy builds on the spectrum-centric DIA approach. It considers all precursors within the possible DDA isolation window and performs a database search. Precursor extracted ion chromatogram (XIC) detection is only performed after database searching in a targeted manner since the RT and m/z was determined from the top PSM, followed by rescoring and iterating over the potential subsequent PSMs. The similarity in the PSM determination, as well as downstream processing provides and an appropriate framework for an objective DDA and DIA comparison.

### Database searches

All search settings are included in Supplemental table 2. DDA and DIA files were processed using FragPipe (version 23.0) and all files were searched individually, unless specified otherwise. DDA files were searched in DDA+ mode. DIA files were searched using the default SpecLib Quant workflow, with minor changes. In overview, identifications were made using the MSFragger^31–33^ (4.2) search engine followed by validation using MSBooster^34^ (1.3.9) and Percolator^35^ (3.7.1). Protein grouping was performed by ProteinProphet^36^ and FDR filtering by Philosopher^37^ (5.1.1). MS1 quantification was performed using IonQuant^38^ (1.11.9) for DDA+. For DIA files, a spectral library was generated using EasyPQP (0.1.52) and quantification performed using DIA-NN^39,40^ (1.8.2_beta_8). Files used for DDA method development found in Supplemental Fig. 1 were acquired and searched over a longer period of time so both FragPipe 21.1 and 22.0 were used. These settings are included in Supplemental table 2.

Before database searching, peak picking was performed within FragPipe for DDA, where the number of peaks used per spectrum was determined automatically, no minimum intensity threshold was applied, and centroiding was verified. DIA files were first converted to mzML format using MSConvert (3.0.23208-1321c92, ProteoWizard^41^) with the vendor peakPicking filter applied. The files were searched against the Human proteome (UP000005640, downloaded 29th April 2024) from Uniprot. The species mix files were searched against the human proteome combined to the Escherichia Coli proteome (strain K12, UP000000625, downloaded 16th May 2025). The FASTA files also contained contaminants and consisted of 50% decoy sequences. MSFragger mass calibration and parameter optimisation was activated, performing a first search and defining the precursor mass tolerance to use for the main search. Protein digestion was set to trypsin allowing for 2 missed cleavages for DDA+ and the default of 1 missed cleavage was left for DIA. Cysteine carbamidomethylation was set as a fixed modification.

Methionine oxidation and N-terminal acetylation were set as variable modifications. A maximum of three variable modifications were allowed per peptide. DDA+ mode was used, with the default setting of 5 top N peptides reported. MSBooster was used for rescoring using the DIA-NN and AlphaPept MS2 Generic models for RT and spectral rescoring, respectively. These models were selected based on the best model test functionality within FragPipe, utilising Koina^42^, applied across multiple datasets, which consistently identified these two models as providing optimal performance for our data. Percolator was used for PSM validation, PTMProphet^43^ for phosphorylation localisations and ProteinProphet for protein inference. DDA+ relied on IonQuant for MS1 label free quantification (LFQ). MaxLFQ values were obtained with a minimum of two ions. For the replicate combined searches, match between runs (MBR) with a 0.01 ion FDR was used and intensity normalisation across runs was enabled. DIA quantification was performed using DIA-NN using a FDR 0.01 cut-off, high precision quantification strategy. For combined file searches, unrelated runs and run specific protein FDR options were not used.

Slight modifications to the standard workflow were made for some specific searches. For the quantitatively matched methods, both DDA+ and DIA utilised “0/1/2” isotope error settings and stricttrypsin with 2 missed cleavages for MSFragger identifications, then in the validation tab, both percolator minimal probabilities were set to 0.5 and FDR filtering and report generation set to “—picked --prot 0.01”.

Finally, for phosphoproteome database search, STY phosphorylation was set as a variable modification. To provide a more direct comparison, the default DIA workflow was modified to resemble the DDA parameters more closely by increasing the number of missed cleavages from one to two and the maximum occurrences of STY phosphorylation per peptide set to 3 in contrast to the standard of 2 in the default DIA_SpecLib_Quant_Phospho workflow.

### Picked Group FDR

The dataset shown in Supplemental Fig. 2 was acquired using thirty replicates. To ensure peptide and protein gains were not linked to false positive accumulation, the data was search using FragPipe without FDR filtering then the picked group FDR strategy^44^ (0.9.0) was applied. The FragPipe search was modified to remove FDR filtering achieved by removing Percolator minimum probability (0.0 vs 0.5 default) and filter section was replaced with “--prot 1.0 --ion 1.0 --pep 1.0 --psm 1.0”. The PSM files from FragPipe containing Target and Decoy assignments were grouped to peptide level. Picked group FDR was run on the grouped files and quantification was disabled. Finally, the updated FragPipe files were filtered to reach 1% FDR.

### Data Analysis

The majority of data was processed in Python (3.12.3) using pandas (2.2.1), numpy (1.26.4). Figures were generated using matplotlib (3.10.0) and seaborn (0.13.2) then adjusted in Inkscape (1.1.2). Picked group FDR was run using Python 3.11.15 and the picked group FDR package (0.9.0). The number of matched ions was extracted from .pepxml files using pyteomics (4.7.2).

Data processing was kept to a minimum to allow close examination of the underlying data. No data points were removed or imputed. For protein identifications, only proteins supported by at least two peptides, including a minimum of one unique peptide, were retained to ensure reporting confidence. For intensity comparisons, MS1 and MS2 intensities were log2-transformed and median-centred prior to visualisation in density plots. For certain comparisons (specified in main text and figure captions), only peptides common to both DDA and DIA were included.

Phosphopeptide localisation was handled as follows. For each peptide, only the localisation with the highest probability score was retained. For multiply phosphorylated peptides, where multiple sites could be assigned, the number of retained localisations was matched to the number of phosphorylated residues per peptide, such that each phosphorylation was assigned to its most probable site. This approach ensured that no peptides retained ambiguous localisation assignments. We acknowledge this simplification may not reflect best practices for biological studies, where localisation confidence thresholds are typically applied. Here, no such thresholds were used, as the goal was to examine technical differences between acquisition methods rather than draw biological conclusions from individual phosphorylation sites. A small proportion of DDA phosphopeptides lacked a localisation probability entirely, likely due to low precursor intensity; these peptides could not be included in localisation analyses and represented 4% of all DDA-identified phosphopeptides.

## Results

### Data acquisition speed drives DDA performance for peptide and protein identification

All experiments were performed on an Orbitrap Astral mass spectrometer. To enable a systematic comparison between DDA and DIA, we evaluated a range of DDA MS1 parameters, MS2 filters and MS2 parameters using bulk trypsin digested HeLa cell line lysates and a 22 min active gradient LC method (Figure 1A). Of the MS1 parameters, short maximal injection times (maxIT; Supplemental Fig. 1A) and high resolution (Supplemental Fig. 1B) had positive impacts on peptide identifications while precursor scan ranges had little effect (Supplemental Fig. 1C). Of MS2 filters, a dynamic exclusion duration of five seconds showed best results using our LC setup. This value will vary depending on chromatographic peak width (Supplemental Figure 1D). Changing monoisotopic precursor selection (MIPS) and precursor ion charge filters had little effect (Supplemental Fig. 1E), while using Precursor Fit and Intensity filters should generally be avoided (Supplemental Fig. 1F-G). Concerning MS2 parameters, isolation windows of 1.2 Th and 2.1 Th provided the highest peptide identifications (Supplemental Fig. 1H), which is why we chose the 1.2 Th setting for DDA throughout this manuscript to minimise peptide co-isolation. MS2 maxIT in isolation had little effect on identifications within the narrow range tested (2-5 ms; Supplemental Fig. 1I).

To illustrate the overall effects of the above, we compared the “Optimised” method to an “Exploris-like” method that used typical MS1 and MS2 parameters and filters for this prior-generation instrument as well as a “Fast” method in which all DDA parameters were chosen to achieve highest data acquisition speed (Figure 1B-G; Table 1; Supplemental Table 1). The use of optimised parameters and filters increased identifications by 17% for peptides (Fig. 1B) and 10% for proteins (Fig. 1C). Maximizing speed, provided an additional benefit of 6% for peptides and 3% for proteins despite marginally lower median peptide MS1 intensity (Fig. 1D). These improvements were rooted in the ability to acquire more MS2 spectra during the ∼25 min productive MS time (average of 122,785; 150,325 and 177,236 MS2 spectra for the Exploris-like, Optimised and Fast methods respectively). This resulted in a concomitant total increase of cumulative peptide spectrum match (PSM) counts of 56% (from 152,107 to 209,267 to 237,929 PSMs respectively; Fig. 1E). The Optimised method showed a slightly better conversion rate from MS2 spectra to PSMs compared to the Fast method (139% vs 137%; Fig. 1F), presumably because of better quality MS2 spectra by virtue of longer MS2 ion accumulation times. As a result, the Optimised method performed slightly better for very low sample loading (Fig. 1G).

**Table 1.**
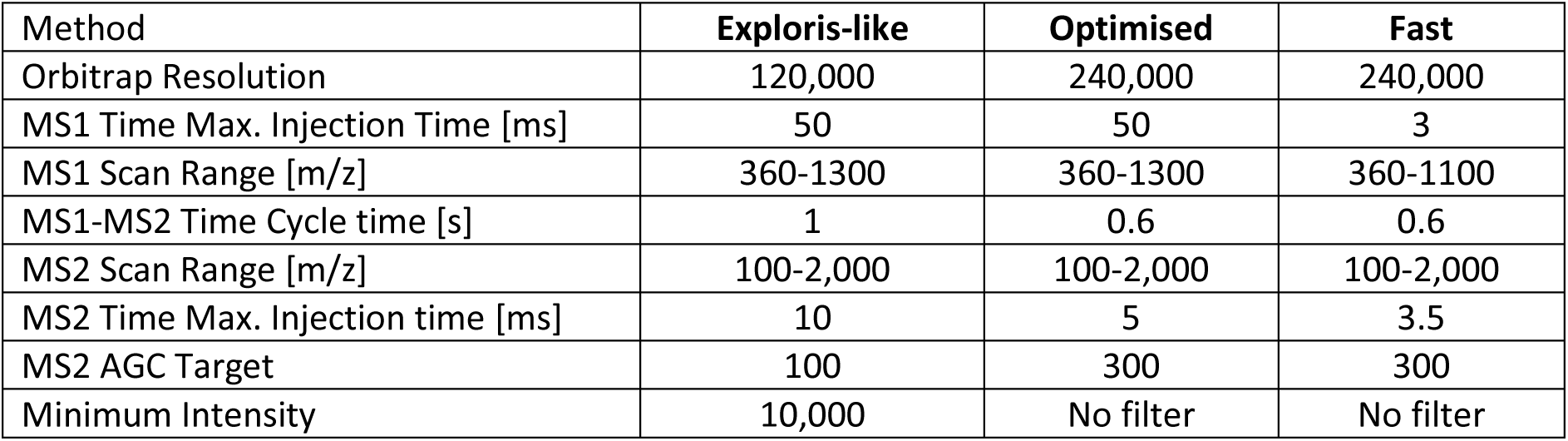
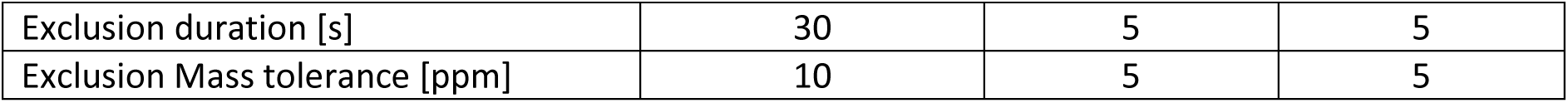
Summary of key parameters for DDA methods parameters.

| Method | Exploris-like | Optimised | Fast |
| --- | --- | --- | --- |
| Orbitrap Resolution | 120,000 | 240,000 | 240,000 |
| MS1 Time Max. Injection Time [ms] | 50 | 50 | 3 |
| MS1 Scan Range [m/z] | 360-1300 | 360-1300 | 360-1100 |
| MS1-MS2 Time Cycle time [s] | 1 | 0.6 | 0.6 |
| MS2 Scan Range [m/z] | 100-2,000 | 100-2,000 | 100-2,000 |
| MS2 Time Max. Injection time [ms] | 10 | 5 | 3.5 |
| MS2 AGC Target | 100 | 300 | 300 |
| Minimum Intensity | 10,000 | No filter | No filter |
| Exclusion duration [s] | 30 | 5 | 5 |
| Exclusion Mass tolerance [ppm] | 10 | 5 | 5 |

### nDIA modestly outperforms DDA for peptide and protein identification and data completeness

It has been shown before that narrow DIA window sizes improve peptide and protein identification results of DIA measurements. One prospect of the high acquisition speed of the Orbitrap Astral or other such mass spectrometers is that the empirically observed differences between DIA and DDA might diminish when using very narrow windows (nDIA) of e. g. 2 Th. To compare DDA and nDIA for peptide and protein identification in more detail, Fragpipe^29,31,32^ was chosen for data analysis because PSM determination between both data types is conceptually similar. Replicate measurements (n=3) using low (1 ng), medium (100 ng) and high sample input amounts (1,000 ng) were acquired using the Fast DDA method described above (1.2 Th selection window) as well as an nDIA method (2 Th window) at a speed of 180 Hz (MS2 maxIT 3 ms). For the sake of being conservative, ≥2 peptides were required for protein identification, of which one peptide had to be unique. In concordance with previous results, nDIA mean peptide (Fig. 2A) and protein identifications (Fig. 2B) were higher for nDIA for all inputs, with the largest difference observed (+35% peptides, +32% proteins) for the 1 ng measurements (+12%/19% for 100 ng input and +8%/16% for 1,000 ng input). All data analysis discussed below was performed using the 100 ng input samples. For both, DDA and DIA, Fragpipe used a median of 7 fragment ions to call a peptide identification common to both methods (Fig. 2C), indicating that the identification algorithms behave broadly similarly with respect to MS2 spectral processing. The MS1 peptide intensity distribution of peptides commonly identified by DDA or nDIA spanned >15 logs and peptides identified exclusively by DDA nearly followed the same intensity distribution. In contrast, peptides identified exclusively by DIA were generally of lower intensity, consistent with the precursor ion-agnostic acquisition of DIA data which benefits the detection of lower-abundance peptides (Fig. 2D).

**Figure 2.**
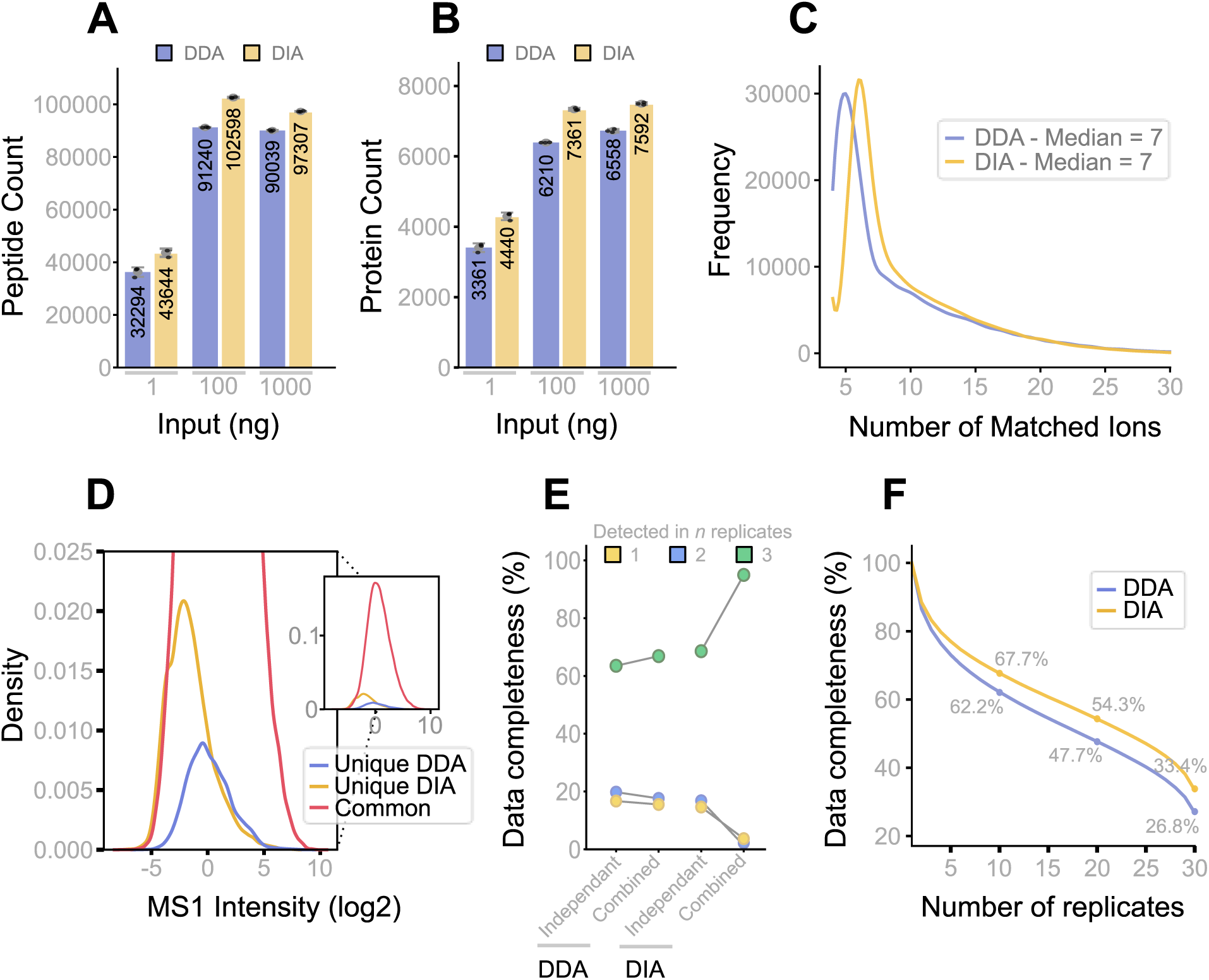
Performance comparison of DDA and nDIA acquisition methods for peptide and protein identification as well as data completeness. (A) Bar plots showing the number of identified peptides for three sample input amounts of HeLa tryptic digests (mean +/- standard deviation, n=3) for DDA and nDIA. (B) Same as panel (A) but for proteins. Protein identifications required two or more peptides and a minimum of one unique peptide. (C) Line plot comparing the number of matched fragment ions per peptide identification common to both DDA and nDIA methods using 100 ng Hela tryptic digest on column, the data shown is from replicate 1. (D) Zoomed density plot showing the distributions of MS1 intensities of identified peptides. Peptides unique to DDA are shown in blue, peptides unique to nDIA are shown in yellow. Peptides identified in both are shown in red. The inset shows the complete density plot. (E) Proportion of peptides identified in one, two, or three of three DDA and nDIA replicates, comparing independent database searches of individual replicates with a combined database search across replicates. (F) Line plot showing the proportion of peptides identified in at least 1-30 replicates, after independent searches followed by picked group FDR filtering for DDA and nDIA. Peptide presence was calculated for DDA and DIA independently. Data completeness at 10, 20 and 30 replicates are highlighted.

Data completeness between experiments is one of the main conceptual benefits of DIA over DDA measurements because DIA acquisition should not suffer from the semi-stochastic nature of collecting DDA data. However, when searching replicates independently, the difference in data completeness between DDA (64% of peptides identified in 3/3 replicates) and DIA (69%) was rather modest (Fig. 2E). When searching replicates together, data completeness for DDA only increased from 64% to 67%. In stark contrast, the combined DIA searches increased data completeness from 69% to 95%. We suspect that the large difference in the gains in data completeness may be rooted in the process and stringency with which ionquant or DIA-NN assign matches between experiments. IonQuant performs match-between-runs (MBR) by transferring RT-aligned precursor features between DDA runs using tight m/z and RT tolerances consistent with those observed in independently identified runs. In contrast, DIA inherently uses a shared library leveraging identifications from all runs. Furthermore, the Dia-NN “MBR” option is not truly a matching algorithm but a peptide-centric spectrum library-driven re-analysis that uses globally aligned retention times and fragment-ion evidence to detect peptides in DIA runs where they could not be identified confidently in isolation. The limited gain observed in DDA replicate searches, therefore, likely reflects very stringent requirements imposed on identification transfers by IonQuant. It has been noted before, that DIA “matching” approaches based on spectral libraries can lead to be overly optimistic estimates of data completeness^30^.

To investigate this further, we expanded the above analysis to 30 identical replicates which led to several surprises. While the mean number of peptides and proteins identified by nDIA were somewhat higher than those for DDA (6% and 5% respectively), the total numbers of unique peptides identified was notably close (166,845 peptides for DDA and 161,255 for DIA). The small gap at the peptide level closed almost completely (1,411 peptides or 0.8%) when removing 6-amino acid peptide identifications from the DDA data (these are categorically excluded for DIA data in Fragpipe).

In addition, for both DDA and nDIA, the number of unique peptides never reached a plateau (despite rigorous FDR, and protein grouping control, see methods; Supplemental Fig. 2C). This suggests that the molecular complexity of the digest was too high to be fully sampled by only a few measurements. In fact, the data suggests that sampling is semi-stochastic for both DDA and nDIA. Contributing factors likely relate to many if not all steps in the LC-MS sequence of events, including small fluctuations in peptide elution from the LC system or electrospray ionization both of which may make a peptide detectable in one LC-MS run but not in another. Despite the overall similar analytical performance of DDA and nDIA for peptide identification, the overlap of all unique peptides detected by both methods was only 69% with the proportion of peptides unique to DDA (17%) or nDIA (14%) being similar (Supplemental Fig. 2E left). Again, when removing peptides of 6 amino acids in length, the non-overlap only marginally decreases in absolute terms. At the protein level, the number of identifications also continued to rise with increasing replicates, albeit at a lesser rate and the difference between protein identifications between nDIA and DDA was below 3% 7,769 for nDIA and 7,563 for DDA, Supplemental Fig. 2D). Here, the non-overlap between the methods was only 8%, with 202 (2.5%) proteins unique to DDA and 408 (5.1%) unique to nDIA (Supplemental Fig. 2E right).

We could not find clear reasons for the large extent of non-overlap on peptide level between DDA and nDIA (Supplemental Fig. 2F-H). The total number of peptides per protein identification were similar, as well as the number of unique peptides per protein. Using GRAVY (hydrophobicity) scores as a proxy for chromatographic behaviour, peptides unique to DDA or nDIA showed essentially the same distribution. There was a notable difference in peptide length and m/z. Peptides exclusively identified by DDA >1,000 m/z were not systematically not measured in nDIA and peptides <7 amino acids were not scored by Fragpipe in nDIA. However, these two factors do not nearly explain the large number of peptides unique to DDA. Again, DIA-only peptides were typically identified with lower peptide abundance, observed both on MS1-intensity level and MS2-intensity level (Supplemental Fig. 2J-K), while DDA-only peptides followed the MS1 intensity distribution of the shared peptides. One plausible reason for nDIA missing many peptides that were indeed detected by DDA, is that these required different collision energies than applied during the nDIA measurement. Likewise, DDA may have missed many peptides detected by nDIA because of too low abundance in a given LC-MS run. The number of total and unique peptides contributing to protein identifications were comparable between DDA and nDIA, suggesting that both DDA-unique and nDIA-unique proteins were genuinely identified rather than the mere result of false discoveries.

Returning to the aspect of data completeness, the 30-replicate analysis showed that data completeness decreased steadily with increasing replicates because of the aforementioned observation that the cumulative number of identified peptides never saturates. Similar trends were observed for both methods (Fig. 2F) with nDIA always more data complete than DDA for any number of replicates but the difference never exceeded 6.6%.

### DDA and nDIA yield similar quantitative precision and accuracy

Replicate analysis (n=3) of 100 ng HeLa digests measured on a 22 min active gradient resulted in an calculated average of 15 MS1 data points per LC peak for DDA, 18.5 MS1 data points for nDIA and 5.8 MS2 data points for nDIA. Analysing the data for quantitative precision showed that 61-66 % of all peptides showed a coefficients of variation (CV) below 20 % (Fig. 3A-B). We note that despite defining a fixed 0.6 second cycle time for both collecting MS1 spectra in DDA and nDIA, differences in ion handling and routing led to differences in actual cycle time in practice (0.68 sec for DDA, 0.55 for DIA). We further note that this analysis was performed on the “raw” intensities of each LC-MS run: each file was searched independently to assess the reliability of a single acquisition and to avoid that CVs are improved by cross-run normalization performed by software. From this data, we conclude that all three approaches yield highly consistent and repeatable quantitative data for this particular 22 min active gradient LC-MS method. To address quantitative accuracy at the peptide level, we analysed controlled mixtures of HeLa and E. Coli peptides (Fig 3C). All three approaches broadly recapitulated the expected log2 ratios of 1.

**Figure 3.**
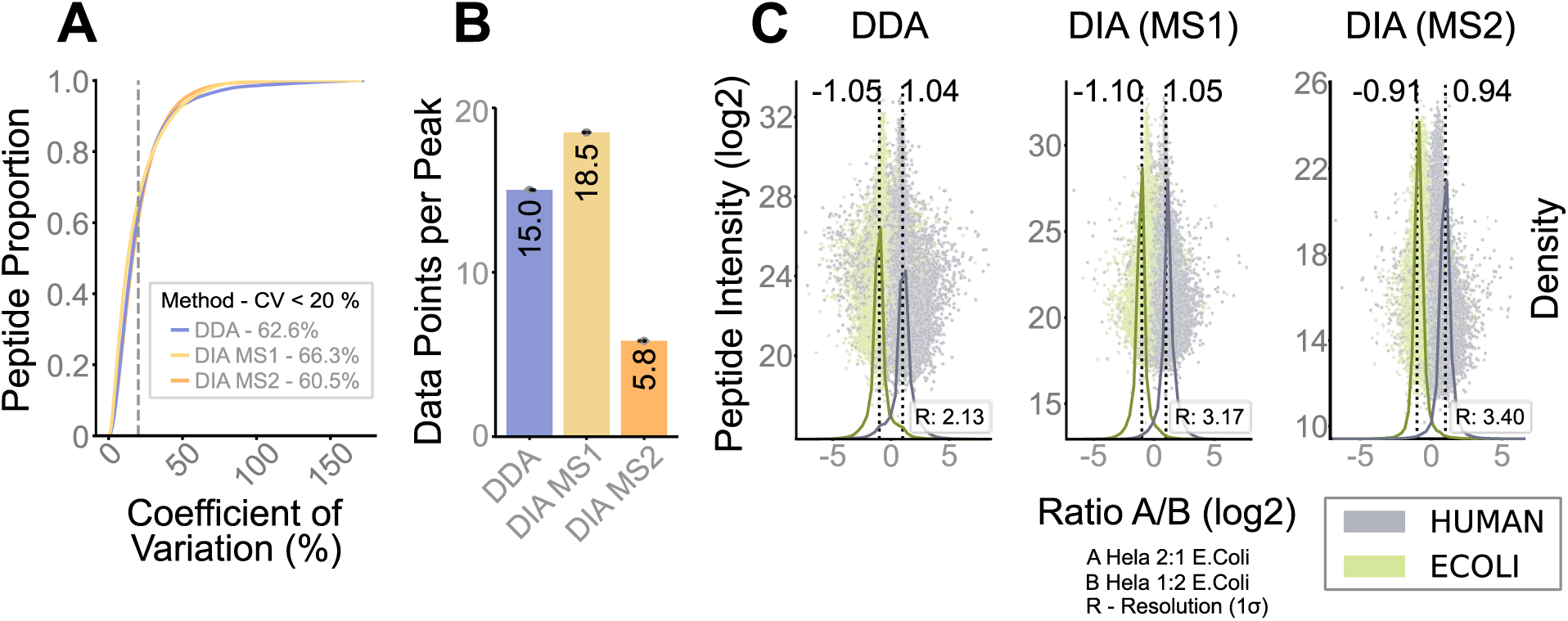
Quantitative performance comparison of DDA and nDIA acquisition methods at the peptide level. (A) Cumulative density plot showing the proportion of identified peptides as a function of quantitative precision (CV: coefficient of variation; n=3, 100 ng HeLa tryptic digest on column). The vertical dashed line marks 20 % CV. (B) Bar plot showing the calculated number of MS1 points derived from the average chromatographic peak width of peptides and scan acquisition frequency for DDA and nDIA as well as MS2 for nDIA (mean +/- standard deviation, n=3). (C) Scatter plot showing the intensity (mean, n=3) of identified peptides derived from E. coli or human sources and the deviation from expected HeLa:E. coli ratios of 2:1 (Ratio A) or 1:2 (Ratio B). Peptide intensities were derived from MS1 signals for DDA (left panel), as well as from MS1 (middle panel) and MS2 intensities (right panel) for nDIA. The scatter plots are overlaid with density distributions (solid lines) for both organisms (HeLa, grey; E. coli, green). Expected ratios (−1 and +1) are indicated by vertical dashed lines, and observed ratios (density peak apex) are reported above each distribution. The resolution (R:) by which the organism density lines are separated is shown and was calculated using R = peak apex Human – peak apex Ecoli / 0.5 (σ human + σ Ecoli; σ being the standard deviation of the respective distributions).

Global DDA and nDIA MS1 ratios were slightly expanded whereas MS2 ratios were compressed. Global accuracy was marginally better for DDA than for nDIA using MS1 intensities and both were more accurate than using MS2 intensities of the nDIA data (average absolute errors of the ratios (log2) of 0.045 for DDA and 0.075 for DIA (both MS1 and MS2 levels)). We also calculated the resolution of the ratio densities, which can be viewed as a measure of precision. Here, nDIA (MS2) performed best, followed by nDIA (MS1) and DDA. The higher resolution of the nDIA MS2 data can be rationalised by the (mostly) absence of interfering signals, which are more likely to occur in MS1 spectra. The reason why nDIA (MS1) outperformed DDA (MS1) is likely because of ∼20% more data points per LC peak noted above.

At the protein level, DDA-based quantification showed better accuracy than nDIA (average absolute errors of the ratios (log2) of 0.035 for DDA and 0.080 for DIA; (Supplemental Fig. 3)). This difference is likely due to differences in how Fragpipe rolls up peptide quantification into protein quantification. For DDA data, the MaxLFQ algorithm is used, while DIA-NN uses the MS2 data for this purpose. Collectively, these results suggest that, despite methodological differences in data acquisition, both DDA and nDIA deliver good quantitative precision and accuracy at the peptide and protein level with small advantages in terms of accuracy for DDA and precision for nDIA.

### MS1 quantification outperforms MS2 in DDA and nDIA at any LC gradient

The speed of the Orbitrap Astral decreases the measurement time needed per sample to reach substantial proteome coverage, hence improving throughput and examples of LC separations as short as 5 and 7 minutes have been published ^16,17^. Such rapid separations challenge the cycle time needed for nDIA quantification employing the traditional MS2-based approach. To investigate quantitative precision based on MS1 and MS2 data in this context, we analysed HeLa digests using 5-, 15-, 30 and 60- minute active gradients (Fig. 4). First, we measured PROCAL peptide standards^45^ to determine the average chromatographic full peak width of each gradient which ranged from 4.0 seconds (5 min gradient) to 20.2 seconds (60 min; Fig. 4A). The relatively broad peaks observed at 60 minutes were expected given the use of a short Aurora Rapid column (8cm x 75 um) that is typically used for short gradients in high-throughput applications.

**Figure 4.**
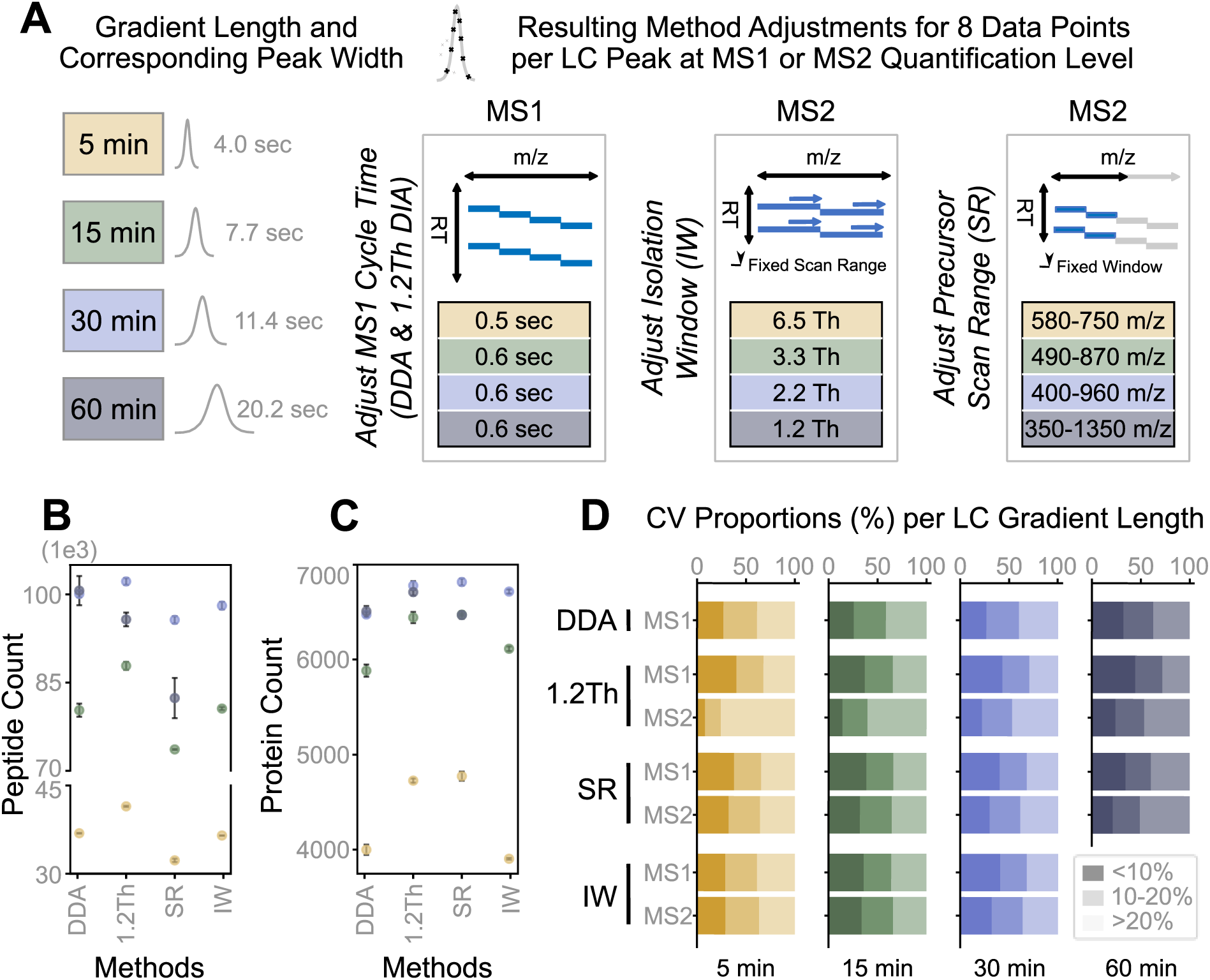
Performance comparison of DDA and DIA methods matched for 8 data points across the LC peak across chromatographic gradients. (A) Schematic overview of the strategy used to achieve an average of ∼8 data points per chromatographic peak across gradients of increasing length (5, 15, 30, and 60 min). Left panel: LC peak widths for different gradient times. Matching colours are used for each gradient throughout the figure (5 min, yellow; 15 min, green; 30 min, blue; 60 min, dark blue). Right panel: Data acquisition parameter adjustments needed to fulfil the 8 data point criterion: optimisation of MS1 cycle time (left), MS2 isolation window width (middle), and precursor scan range (right). (B) Number of peptides identified for each LC gradient and data acquisition method (mean +/- standard deviation, n=3). (C) Same as panel (B) but for proteins. Protein identifications required two or more peptides and a minimum of one unique peptide. (D) Stacked bar plots showing the proportions of peptides quantified within three ranges of precision for each LC gradient (shaded colours; CV: coefficient of variation).

Guided by published work^46^, we selected an average of eight data points per LC peak across the gradient for all experiments below to ensure comparability and adequate quantitative performance. Fixing the desired number of data points per LC peak as a parameter requires using adequate cycle times of the mass spectrometer. For MS1-based quantification in DDA and nDIA (both using a 1.2 Th selection window and 360-1,000 m/z MS1 scan range), this is readily achieved owing to the high degree of parallelization between MS1 and MS2 scans on the Orbitrap Astral. For the 5’ gradient, a cycle time of 0.5 seconds and an Orbitrap resolution of 180,000 met the 8 data point criterion. A cycle time of 0.6 seconds enabled operating the Orbitrap at 240,000 resolution and still provided sufficient sampling for gradients of 15 min and above (Fig. 4A). For MS2-based quantification (only applicable to DIA), adequate sampling of the LC peak can be achieved in two ways, with a mix of both often applied in practice: i) adapting the precursor ion isolation window to cover a desired MS1 m/z scan range (IW method; here 360-1000 m/z) or ii) restricting the MS1 m/z scan range to maintain narrow precursor ion isolation (SR method; here 2 Th). For the IW approach, this translated into isolation windows of between 6.5 and 1.2 Th. For the SR approach, MS1 scan ranges 170 – 1,000 m/z wide could be covered in a single LC-MS run.

The data of these comparisons are summarized in Fig. 4B-D and reveal a few patterns. First, the differences in terms of peptide and protein identification were not dramatic (Fig. 4B,C) but DDA (1.2 Th) and ultra nDIA (1.2 Th) provided the highest number of peptides confirming that narrow isolation windows benefit DIA identifications by generating MS2 spectra of low complexity. As expected, the overlap of peptides identified by DDA and nDIA increased from 47% to 77% for the 5 min and 60 min LC gradients respectively (Supplemental Fig. 4A). The SR method performed worst at short gradients because only a small m/z range of the peptidome could be covered. Second, all methods performed best in absolute terms using a 30 min gradient, supporting prior conclusions for DIA and here confirming for DDA that the Orbitrap Astral is not substantially more productive at extended LC gradients^17^. This is likely due to the broader LC peaks at 60 min which reduces peptide concentration and sensitivity. We note that the use of other column materials or flow rates will have an influence on this assessment.

Third, at the protein level (Fig. 4C), all DIA methods modestly outperformed DDA. Interestingly, the SR method compensated the lower peptide coverage by a gas-phase fractionation effect which led to protein identifications that were on par with that of the ultra nDIA method.

In regards to quantitative precision, MS1-based measurements were consistently more repeatable than MS2-based measurements at any gradient time (Fig. 4D). MS1 cycle times were again deliberately set to a minimum of 8 data points per peak but many measurements actually exceed this threshold, potentially explaining the observed differences. Also in this metric, DIA MS1 outperformed DDA MS1 which may be attributed to the slightly higher MS1 cycle time in DDA in comparison to DIA (see Fig. 3B above). In addition, the proportion of peptides with CVs <20% improved with longer gradient times which was also true for peptides found in all measurements (Supplemental Fig. 4.B). The poorest CVs were found for MS2 quantification in the 1.2 Th nDIA method on the shorter gradients (5- and 15-minutes). However, this was expected, as this method was never intended to provide adequate MS2 quantification. All things considered, the data suggests that ultra-narrow-window DIA acquisition with MS1-based quantification provides a strong balance between proteome coverage, quantitative precision and sample throughput.

#### Ultra narrow isolation window DIA closes the gap to DDA performance for phosphoproteomics

Despite the wide adoption of DIA in proteomics, one notion in the field is that DDA is better for phosphoproteomics. In principle, (ultra-)narrow isolation DIA windows that are the same as the ones used in DDA should mitigate challenges associated with the expanded search space in phosphoproteome analysis while retaining the systematic sampling advantage of DIA. Recent studies employing 2 Th and 4 Th DIA methods^18,47^ do point into this direction. Here we expanded this analysis to include nDIA measurements employing 1.2 Th, 2 Th and 4 Th windows with a small restriction on MS1 scan range (360-1000 m/z) to enable adequate quantification and compared these to DDA using a 1.2 Th isolation window (360-1300 m/z scan range, no restriction required).

Replicate analysis (n=3) showed that DDA, 1.2 Th nDIA and 2 Th nDIA performed almost identically (phosphopeptide input of 200ng; 22-minute active gradient) but a substantial drop was observed for the 4 Th nDIA method (Fig. 5A). Focussing on the DDA and 1.2 Th nDIA method showed that the overlap of phosphopeptides between replicates was slightly higher for DDA (45%) than for DIA (41%) but these figures were substantially lower than those observed for non-phosphorylated peptides (Supplemental Fig. 5A-B). The overall quality of phosphopeptide spectra was not different to those of non-phosphorylated peptides judged by the number of matched fragment ions (median of 8-9 for phosphopeptides; Fig. 5C and a median of 7 for non-phosphorylated peptides; Fig. 2C). Both datasets together identified 45,338 phosphopeptides but only 50% were the same (Fig. 5B). This suggests again that the diverse population of phosphopeptides present in the sample overwhelms the analytical capacity of the LC-MS system. As a result, sampling by both DDA and nDIA has a substantial stochastic component. In terms of quantitative precision, phosphopeptides followed the characteristics observed for non-phosphorylated peptides in that 65-73% of all phosphopeptides showed CVs below 20 % when using MS1-based intensities (Fig. 5D). Also, here nDIA (1.2 Th) showed the best overall performance.

**Figure 5.**
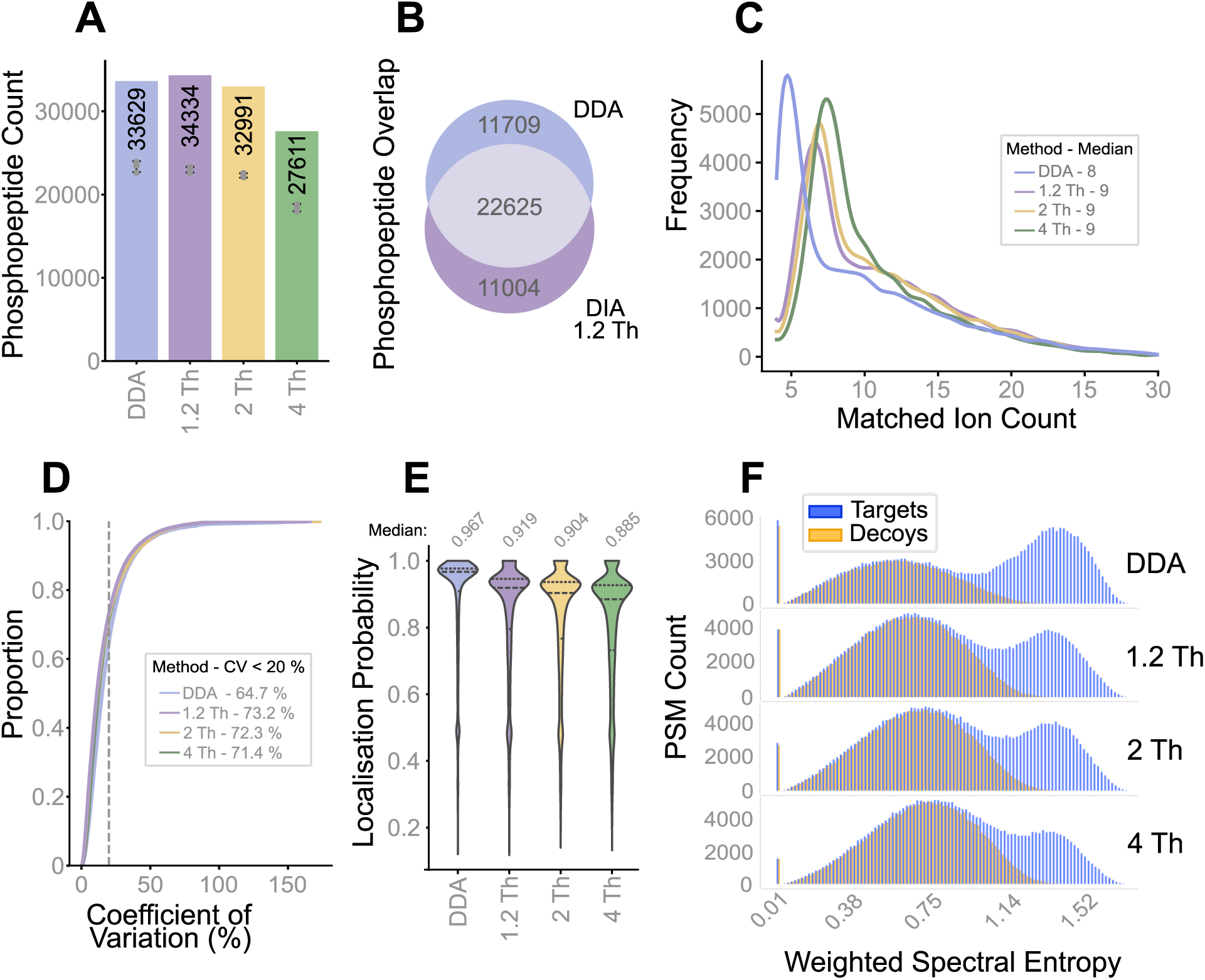
Comparison of DDA and DIA acquisition methods for phosphoproteome analysis. (A) Bar plot showing the total number of phosphopeptides identified using DDA (1.2 Th isolation window) and DIA methods with isolation windows of 1.2 Th, 2 Th, and 4 Th (200 ng phosphopeptide on column). Replicate values (dots, n=3) and standard deviations (bars) are shown in grey. (B) Venn diagram showing the overlap of phosphopeptide identifications between 1.2 Th DDA and 1.2 Th DIA methods. (C) Line plot showing the number of matched fragment ions per phosphopeptide identification for phosphopeptides identified by all methods. (D) Line plot showing the cumulative proportion of phosphopeptides quantified using MS1 data as a function of quantitative precision (CV: coefficient of variation). The vertical dashed line indicates 20% CV. (E) Violin plots illustrating phosphosite localisation probability distributions for DDA and nDIA methods. (F) Distributions of target and decoy peptide-spectrum matches (PSMs) based on spectral entropy distributions (MSBooster) for DDA and nDIA acquisition methods.

An important additional challenge in phosphoproteome analysis is the localisation of the phosphorylation site. Here, DDA was leading with a median localisation probability of 0.97. The median for DIA decreased as isolation window width increased (Fig. 5E) from 0.92 (1.2 Th) to 0.68 (4 Th). Phosphopeptides common to DDA and nDIA (1.2 Th) showed very similar localisation scores (0.97 for DDA, 0.91 for nDIA) but decreased substantially for those unique to either method (0.86 for DDA, 0.77 for nDIA; Supplemental Fig. 5C). The higher performance of the DDA data could not be rationalised by i) how many fragment ions were matched by the search algorithm, ii) how many amino acid residues separated two possible phosphorylation sites, iii) the distance of the phosphorylation site from either the N- or C-terminus of the peptide, iv) the precursor ion charge state, v) the number of missed tryptic cleavage sites or vi) the ratio of singly vs. multiply phosphorylated peptides (Supplemental Fig. 5D-H).

What made a clear difference between the DDA and DIA data was that the target and decoy distributions were better separated in DDA than nDIA (Fig. 5F). Fragpipe contains the processing tool MSBooster^34^ that extracts peptide sequences from search results and feeds them into deep learning architectures to predict features such as RT, and MS2 fragment ion spectra. Among other metrics, it utilises spectral entropy which provides a measure of spectral complexity and confidence in identifications^48^. Higher values are associated with more confident identifications, whereas lower entropy values come with an increased likelihood of random peak matches. Although not the focus of this study and somewhat speculative, a number of possible reasons for the poorer separation of target-decoy distributions in nDIA come to mind: 1. MSBooster uses a single collision energy for an LC-MS run, hence treats precursors of different charge states the same even though their optimal collision energies are different. 2. It is possible that predictions for DDA fragment ion spectra are better than those for DIA data which would likely result in differences in target-decoy separations. 3. DDA search engines can make use of all fragment ions in an MS2 spectrum, whereas DIA search engines often ignore low m/z y-or b-type fragments because they can be found in many peptides. For instance, tryptic peptides almost always have a Lys or Arg at the C-terminus, hence the y1 ion is not very informative unless the expected (i.e. predicted) intensity is taken into account^30^. A similar argument applies to b2 ions. 4. A further contributor could be that DIA is ignorant to selecting peptides based on a detected precursor ion isotopolog. Therefore, many nDIA MS2 spectra will contain “the wrong” isotopes. Deconvoluting these cases may not always work without error.

## Discussion

Each technological leap in mass spectrometry is inevitably followed by benchmarking studies aimed at redefining the boundaries of what is analytically achievable. The current work aimed at providing such a technical benchmark for DDA and DIA analysis on an Orbitrap Astral platform and confirmed many empirical observations made by practitioners in the field. At the same time, the study also highlights a few important limitations and open issues that are sometimes overlooked or ignored. As a disclaimer, we specifically refrained from including a comparison of software for DDA and DIA data analysis in this study for the sake of clarity but acknowledge that this is a limitation.

In light of the results obtained, our main conclusions are as follows:

First, the combined speed and sensitivity of the instrument have enabled the implementation of ultra-narrow precursor ion isolation windows (1.2 Th) for nDIA analysis whose selectivity approach that of a DDA analysis while retaining the advantage of sampling lower abundance peptides. Still, it would be highly desirable if MS instruments would accommodate scan speeds of 500-1,000 Hz to be able to cover the full m/z range of tryptic peptides with an isolation window of 1.2 Th or, conceptually even more powerful, by scanning the quadrupole continuously as recently introduced on the ZenoTOF 7600+ platform.

Second, nDIA data does not appear to be nearly as complete as many reports in the literature imply, at least when taking the analytical data at face value. Our data on 30 identical replicates showed that the number of identified peptides and proteins never saturate reflecting the inability of the LC-MS system to cope with the sheer number of tryptic peptides generated from a proteome. This data set may prove useful to others wishing to investigate the influence of different data processing concepts and software implementations on data completeness in more detail.

Third, turning high-speed mass spectrometers into higher throughput of proteome analysis requires using shorter LC gradients. In our view, the consequence is that quantification in nDIA needs to move away from MS2 to MS1 level data. The results obtained here show that MS1 quantification is robust, precise and accurate and can be generally applied across sample types. Among the reasons for the higher performance of MS1 data is that the Orbitrap has substantially higher ion capacity than the Astral mass analyser, regularly accommodating 10^5^ ions while maintaining linear and accurate quantification, whereas the Astral analyser is reported to show space charging effects already at ∼10^3^ charges^49^. While the single-ion detection capabilities of the Astral provides ultimate sensitivity, these signals can also reflect chemical noise or non-peptidic components complicating downstream analysis^14^. The conceptual advantage of using MS1 signal for quantification on the Orbitrap Astral has been leveraged in the context of single cell and low-input proteome analysis^20^ ^23^.

Fourth, DDA is not dead, yet. While nDIA has closed the gap to DDA in terms of the number of phosphopeptides that can be identified, the localisation of the precise amino acid was globally stronger in DDA data. This is likely linked to a more appropriate choice of normalised collision energy (NCE) in DDA measurements. While for DDA, the NCE is chosen based on the m/z and charge state of the detected precursor ion, the choice of NCE in nDIA is based on the centre of the isolation window under the assumption of the presence of doubly charged precursors. The fact that most MS2 spectra are chimeric, this assumption cannot be generally true. For the same reason, the issue also partly applies to DDA data. This may be addressed by applying a stepped NCE strategy, but doing so massively (40-70%) slows down the instrument. This again, calls for even faster MS platforms than routinely available today.

Despite continued and sometimes step-changing improvements in LC-MS technology, current instrumentation remains far from providing comprehensive proteome coverage in a single shot. The sheer number and abundance range of tryptic peptides derived from full proteomes simply overwhelm the analytical capabilities of any current system. The conceptual solutions to this challenge for LC-MS hardware are not new but increasingly difficult to implement. More dynamic range in sampling the electrospray ion beam will be required to detect more peptides. On-line and on-the-fly separation by chromatography, ion mobility, very narrow or sliding isolation windows will all further improve selectivity. All of these require more instrument speed while minimising ion losses and maintaining single ion detection sensitivity inside the mass spectrometer at the same time. Importantly, and although explicitly not covered in this study, data processing software needs to keep up with new capabilities of MS hardware and the data characteristics it generates. In order to sustain high-quality results, the evolution of search and quantification algorithms require full awareness of the analytical capabilities and limitations. This, more than ever, calls for strong cooperation between data producers and data processors so that data consumers can rest assured that reliable measurements support meaningful biological or biomedical interpretation and insight.

## Data Availability

All data, including raw files, FragPipe search results and supplemental combined tables are available on PRIDE (Proteomics IDEntification Database). Annotated spectra can be viewed using the PDV viewer with psm.tsv files and the associated raw files as input.

## Author Contributions

N.O., F.B, B.K. contributed to conceptualisation, investigation, methodology and writing (draft, review and editing); N.O. curated, analysed, visualised and validated the data; B.K, C.M. provided funding and supervision.

## Supplementary data

This article contains supplemental data. Supplemental Figures: Supplemental_figures.pdf

Supplemental Table 1: ST1_MS_methods.csv – contains summarised MS methods. Supplemental Table 2: ST2_Fragpipe_workflows.xlsx – contains all database search information.

Supplemental Table 3: ST3_F1_S1_DDA.xlsx – contains summarised data associated to Figure 1 and Supplemental Figure 1.

Supplemental Table 4: ST4_F2_dilution.xlsx – contains summarised data associated to Figure 2.

Supplemental Table 5: ST5_S2_30_replicates.xlsx – contains summarised data associated to Supplemental Figure 2.

Supplemental Table 6: ST6_F3_S3_quantification.xlsx – contains summarised data associated to Figure 3 and Supplemental Figure 3.

Supplemental Table 7: ST7_F4_S4_gradients.xlsx – contains summarised data associated to Figure 4 and Supplemental Figure 4.

Supplemental Table 8: ST8_F5_S5_phospho.xlsx – contains summarised data associated to Figure 5 and Supplemental Figure 5.

---

**Supplementary Figure S1.**
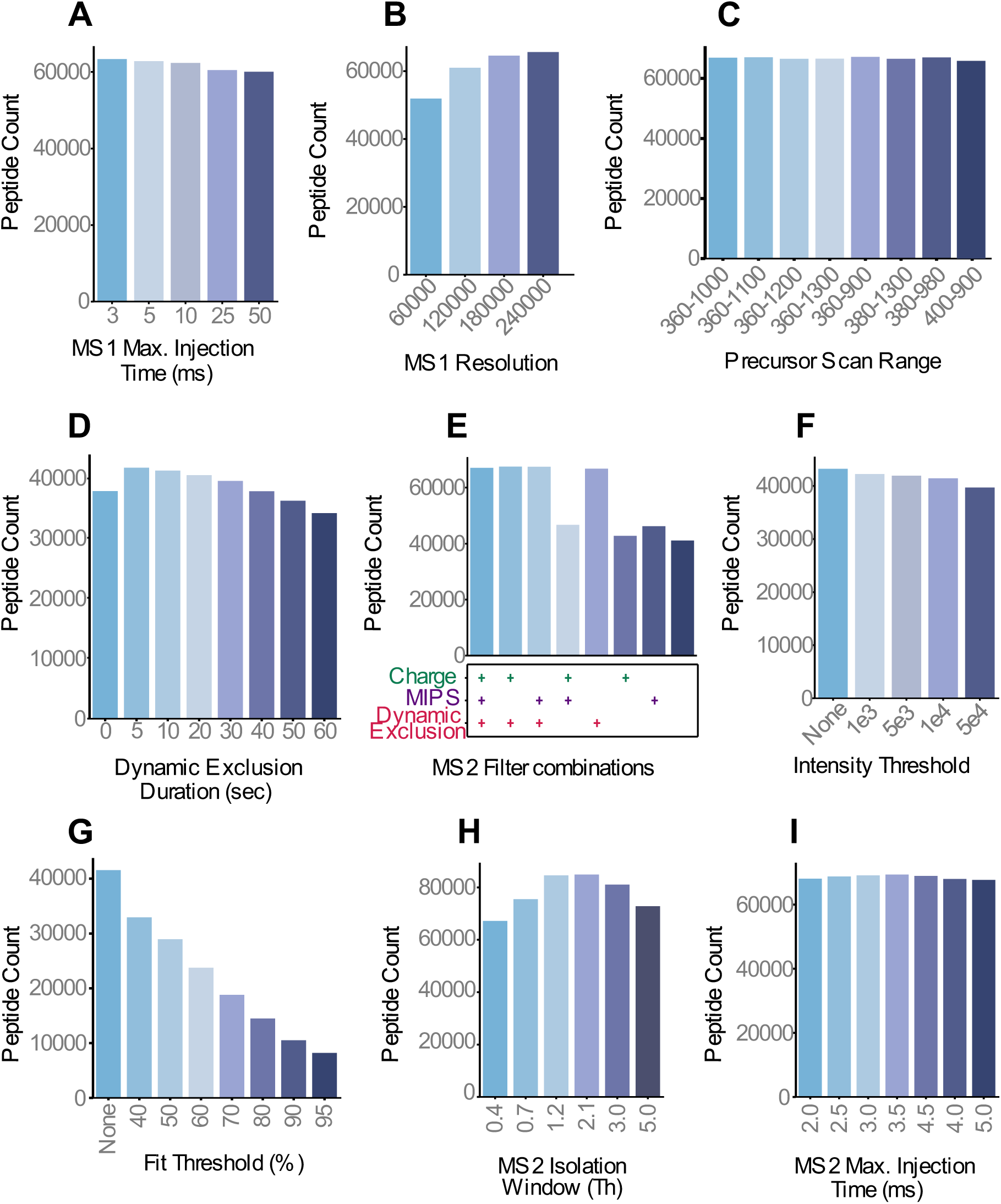
Evaluation of Orbitrap MS1 and MS/MS parameters for peptide identification by DDA. (A) Number of identified peptides (peptide count) as a function of Orbitrap MS1 maximum injection time. (B) Same as panel (A) but for Orbitrap MS1 resolution settings. (C) Same as panel (A) but for Orbitrap MS1 m/z scan range. (D) Same as panel (A) but for chromatographic dynamic exclusion time. (E) Same as panel (A) but as a function of combining different MS2 filter parameters. (F) Same as panel (A) but for different MS1 intensity thresholds for picking precursors for fragmentation. (G) Same as panel (F) but for different precursor ion fit thresholds. (H) Same as panel (F) but for different MS2 isolation windows. (I) Same as panel (F) but for different MS2 maximum injection time settings. Note: performing n=1 measurements for each parameter screening condition was deemed sufficient to observe trends.

**Supplementary Figure S2.**
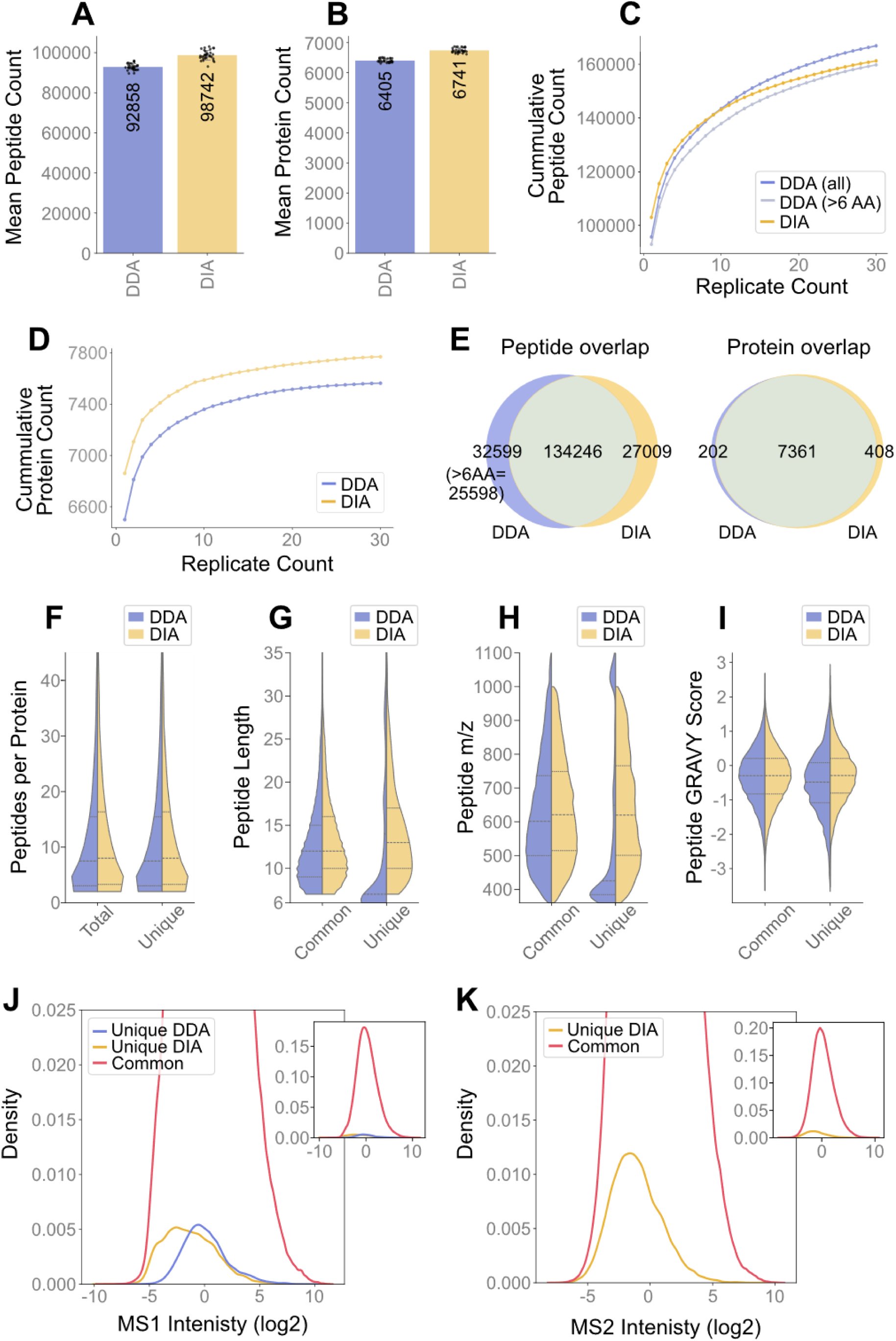
Comparison of DDA and nDIA sampling characteristics across thirty identical replicates. (A) Bar plot comparing the mean number of peptide identifications for DDA and nDIA methods using 200 ng HeLa trypic digest (n=30). All data in this figure were obtained by searching each LC-MS run independently. See methods for false discovery control. (B) Same as (A) but for proteins. Protein identifications required two or more peptides and a minimum of one unique peptide. (C) Line plot showing the accumulation of unique peptide identifications as a function of the number of replicates for DDA (all peptides or filtered for >6 amino acides (AA) per peptide) and nDIA. (D) Same as panel (C) but for proteins. (E) Venn diagrams showing the overlap of peptides (left panel) and proteins (right panel) for DDA and nDIA methods. (F) Violin plots comparing the distributions of the number of peptides identified per protein for DDA and nDIA and contrasting protein identifications based on all peptides or peptides unique to the protein. (G) Violin plots comparing the distributions of the length of peptides identified for DDA and nDIA and contrasting commonly identified peptides and peptides unique to one of the two methods. (H) Same as panel (G) but for peptide mass to charge ratio (m/z). (I) Same as panel (G) but for the GRAVY hydrophobicity score. (J) Line plot showing a zoomed-in section of the MS1 peptide intensity distribution of commonly identified peptides (red), peptides unique for DDA (blue) and nDIA (yellow). The inset shows the full distribution. (K) Same as panel (J) but using MS2 intensity (only applicable to nDIA).

**Supplementary Figure S3.**
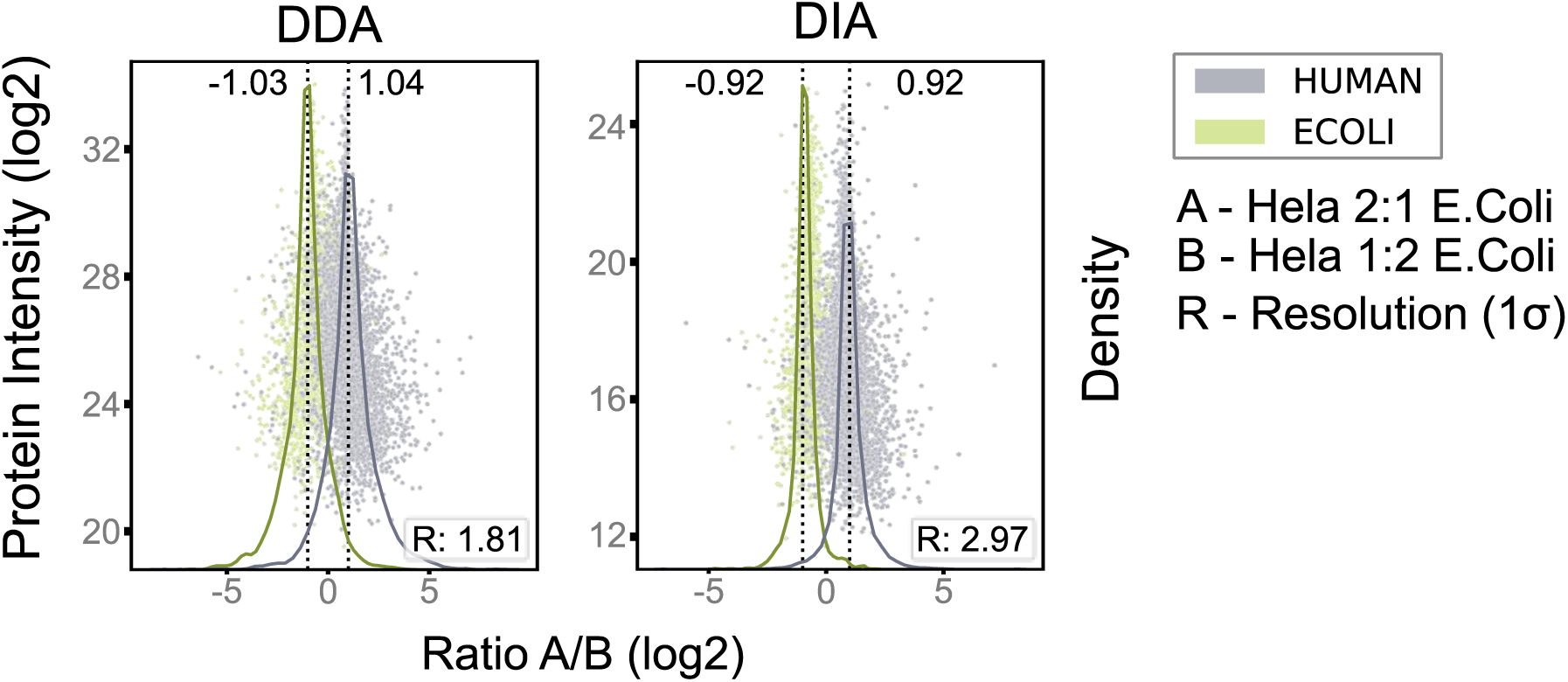
Quantitative performance comparison of DDA and nDIA acquisition methods at the protein level. Scatter plot showing the intensity (mean, n=3) of identified proteins derived from E. coli or human sources and the deviation from expected HeLa:E. coli ratios of 2:1 (Ratio A) or 1:2 (Ratio B). Protein intensities were derived from MS1 signals for DDA (left) and from MS2 signals for nDIA (right). The scatter plots are overlaid with density distributions (solid lines) for both organisms (HeLa, grey; E. coli, green). Expected ratios (−1 and +1) are indicated by vertical dashed lines, and observed ratios (density peak apex) are reported above each distribution. The resolution (R:) by which the organism density lines are separated is shown and was calculated using R = peak apex Human – peak apex Ecoli / 0.5 (σ human + σ Ecoli; σ being the standard deviation of the respective distributions).

**Supplementary Figure S4.**
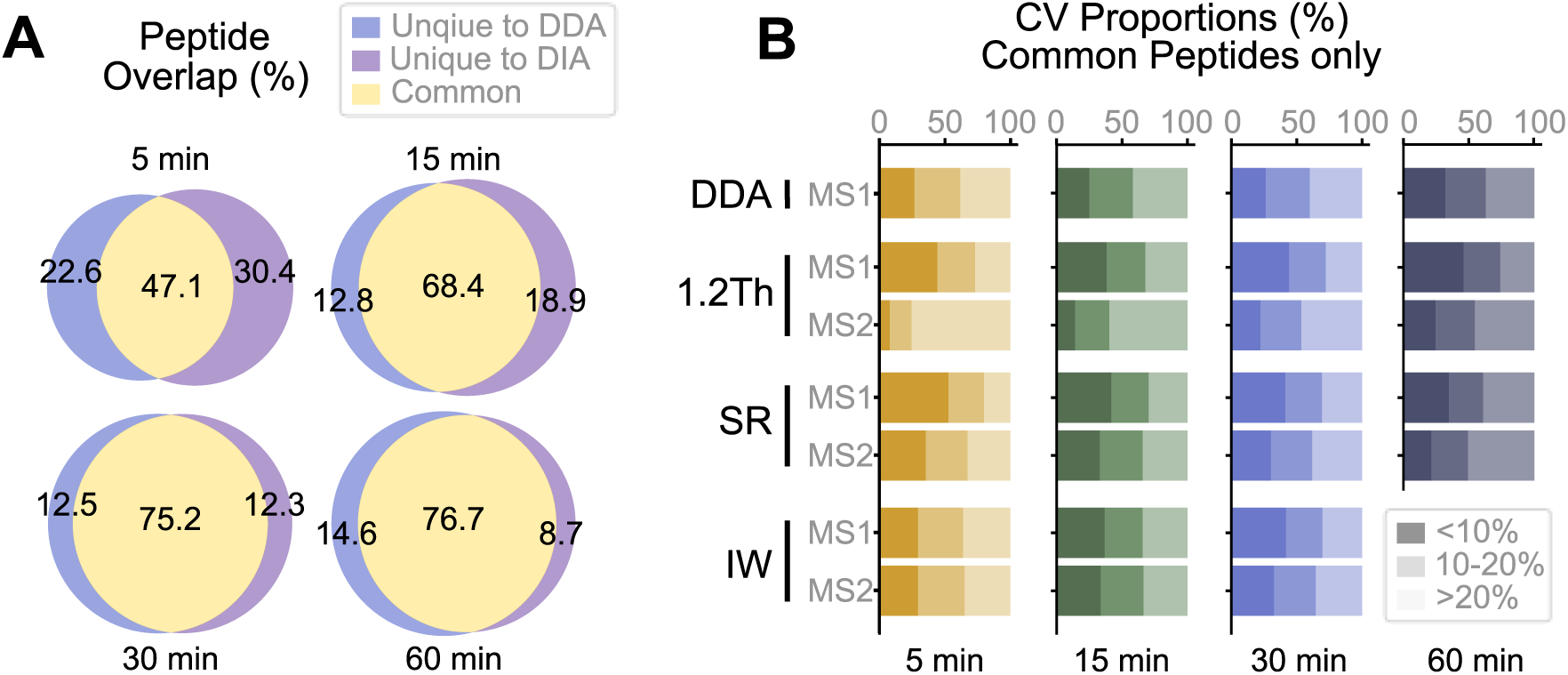
Peptide identification overlap and quantitative precision across chromatographic gradients. (A) Venn diagrams showing the overlap of peptide identifications between DDA and nDIA for each gradient length. (B) Stacked bar plots showing the proportion of peptides within different coefficient of variation (CV) ranges across gradient lengths, calculated for peptides identified by both methods only. Bars are colour-coded by gradient length (5 min, yellow; 15 min, green; 30 min, blue; 60 min, dark blue)., with shading indicating CV ranges (<10%, 10–20%, and >20%).

**Supplementary Figure S5.**
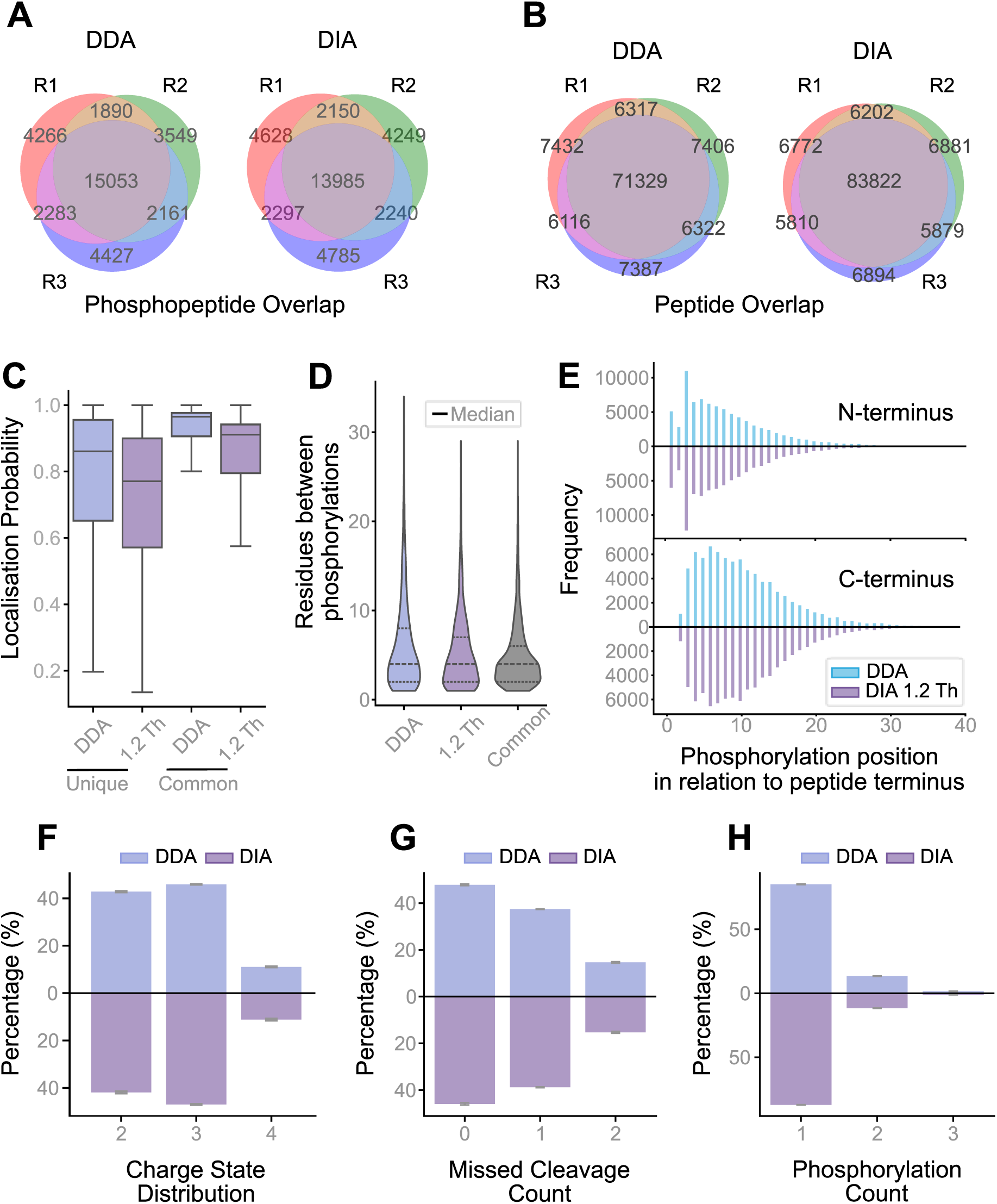
Comparison of DDA and nDIA for phosphoproteome analysis. (A) Venn diagrams showing the overlap of phosphopeptide identifications between three replicates of DDA (left) and 1.2 Th DIA (right) measurements. (B) Venn diagrams showing the overlap of peptide identifications (not enriched) between three replicates of DDA (left) and 1.2 Th DIA (right) measurements. (C) Box plots showing phosphosite localisation probability distributions for DDA and DIA methods, stratified by peptides unique to each method or shared between both. Medians of replicates (n=3) are marked by a horizontal line and whiskers show the interquartile ranges. (D) Violin plots showing the distance (in the number of amino acids) between the phosphorylated amino acid assigned by FragPipe and the next possible phosphorylation acceptor sites within the same peptide. The violins show the distributions of commonly identified phosphopeptides as well as those unique to either method. (E) Mirrored bar plots showing the distance (in the number of amino acids) between the phosphorylated amino acid assigned by FragPipe from DDA and nDIA data and the peptide N-terminus (upper panel) and C-terminus (lower panel). (F) Mirrored bar plots showing the precursor charge state proportions of phosphopeptides identified by DDA and nDIA. (G) Same as panel (F) but for the number of missed trypsin cleavage sites. (H) Same as panel (F) but for the number of phosphorylation events per peptide.

